# Optimal coordination and reorganization of photosynthetic properties in C_4_ grasses

**DOI:** 10.1101/2020.05.15.098327

**Authors:** Haoran Zhou, Erol Akçay, Brent Helliker

**Author notes:** Correspondence:* Haoran Zhou,. Erol Akçay, Brent Helliker.

## Abstract

C_3_ and C_4_ are major functional types in terrestrial biosphere models, with photosynthesis traits as important input parameters. The evolution of C_4_ required reorganizations of Calvin-Benson-cycle and coordination of C_4_-cycle enzymes, resulting in divergences of physiological traits between C_3_ and C_4_. In addition, photosynthesis further optimized after the evolution of C_4_ causing diversification within C_4_ lineages due to different evolutionary histories. We combined optimality modeling, physiological measurements and phylogenetic analysis to examine how various aspects of C_4_ photosynthetic machinery were reorganized and coordinated within C_4_ lineages and as compared to closely-related C_3_ in grasses. Optimality models and measurements indicated a higher maximal electron transport to maximal Rubisco carboxylation ratio (*J*_max_/*V*_cmax_) in C_4_ than C_3_, consistent with the optimal prediction to maximize photosynthesis. The coordination between Calvin-Benson and C_4_ cycles (*V*_pmax_/*V*_cmax_), however, is in line with the optimal modeling results under 200 ppm, as opposed to current CO_2_. Such inconsistencies can be explained by a slowly declining assimilation rate beyond optimal *V*_pmax_/*V*_cmax_. Although rapid coordination occurred early in C_4_ evolution, C_4_ is still under optimizing processes and photosynthetic measures have continued to increase across time. Lastly, better understandings of *J*_max_/*V*_cmax_, *V*_pmax_/*V*_cmax_ and fluorescence-based-electron-transport proffer enhanced approaches to parameterize terrestrial biosphere models.

## Introduction

C_3_ and C_4_ are major photosynthesis pathways and important functional types identified in the terrestrial biosphere models, e.g., Earth System Models (ESMs) and Land Surface Models (LSMs) (Booth et al., 2012; Schaefer et al., 2012; Croft et al., 2017; Rogers et al., 2017). Photosynthetic traits and parameters of C_3_ and C_4_ photosynthesis, e.g., maximal Rubisco carboxylation rate (*V*_cmax_) and maximal electron transport (*J*_max_), are crucial inputs in these models. Such parameters, as well as the *J*_max_/*V*_cmax_, are well-documented for C_3_ photosynthesis, however, the C_4_ photosynthesis are still underrepresented currently due to lack of empirical reports and estimation (Bellasio *et al.*, 2016). The evolution of C_4_ occurred over an extended period that encompassed many different climate change scenarios; therefore C_4_ could respond to the future climate change differently than C_3_ (Ward et al., 1999; Leakey et al., 2009; Wittmer et al., 2010; Morgan et al., 2011; Reich et al., 2018). Additionally, it has recently been proposed that by taking a lineage-based, or evolutionary, approach to ESM and LSM parameterization that a more realistic approach to functional diversity can be captured by these large-scale models (Griffith et al. 2020). Thus, it is crucial to identify the trait divergences of C_4_ photosynthesis from C_3_, especially with regard to the photosynthetic parameters.

C_4_ photosynthesis evolved as a response to inefficiencies of C_3_ photosynthesis that are exacerbated under certain environmental conditions: low CO_2_, drought, high temperature and high light (Ehleringer & Monson, 1993; Ehleringer *et al.*, 1997; Edwards and Smith, 2010; Zhou et al., 2018). Rubisco, the CO_2_ carboxylating enzyme of the Calvin-Benson (CB) cycle also assimilates O_2_ as the first reaction of photorespiration and consequently reduces CB cycle efficiency up to 30% in C_3_ species (Ehleringer *et al.*, 1991; Bauwe *et al.*, 2010; Raines 2011). The C_4_ pathway concentrates CO_2_ around Rubisco and dramatically reduces photorespiration by segregating atmospheric CO_2_ uptake by Phosphoenolpyruvate carboxylase (PEPc) and CB-cycle into two compartments within the leaf. The operation of C_4_ cycle does have additional ATP costs for which C_3_ plants do not remunerate (Hatch, 1987). In sum, the assembly of C_4_ photosynthesis broke the balance between CO_2_ carboxylation and electron transport of the CB cycle, which existed in C_3_ ancestors: C_4_ photosynthesis elevates the efficiency of CO_2_ carboxylation at the expense of using more energy from electron transport (Sage, 2001, 2004, 2016; Christin & Osborne, 2014; Lundgren *et al.*, 2019). Thus, a further reorganization of resource allocation between Rubisco carboxylation and electron transport and the coordination between CB and C_4_ cycle should occur along with, or as a consequence of, C_4_ formation. Furthermore, selection optimized the function of the C_4_ through intermediate steps through adjusting physiological traits (Christin & Osborne, 2014; Sage, 2016) and different C_4_ lineages evolved at different time points and endured different evolutionary histories. We therefore expect diversification in photosynthetic traits and parameters among C_4_ lineages. If C_4_ photosynthesis is continuously under extended optimization after its formation (Edwards, 2019; Heyduk et al., 2019), such diversification would also represent evolutionary trends between photosynthetic parameters and evolutionary time/ages. In the current study, we examine how various aspects of C_4_ photosynthetic machinery— nitrogen allocation between Rubisco carboxylation and electron transport, and CB and C_4_ cycle— were reorganized and coordinated within C_4_ lineage, as well as compared to closely-related C_3_ species, and whether the coordination of C_4_ machineries are on a walk toward or have already reached the optimal state.

The relative ratio among *V*_cmax_, *J*_max_ and maximal PEP carboxylation rate (*V*_pmax_) could represent the coordination within CB cycle and between CB and C_4_ cycles. The resource allocation between the Rubisco carboxylation and electron transport is represented by *J*_max_/ *V*_cmax_. Although *J*_max_/*V*_cmax_ has been empirically measured in numerous C_3_ species (Wullschleger, 1993) and examined with optimal modeling results of *J*_max_/*V*_cmax_ for C_3_ (Walker *et al.*, 2014; Kromdijk & Long, 2016; Quebbeman & Ramirez, 2016), there have been far fewer measurements or predictions in C_4_. The coordination among CB cycle and the C_4_ cycle could be represented by the relationships among *V*_cmax_, *J*_max_ and *V*_pmax_. As a relationship between *J*_max_ and *V*_cmax_ has been established above, *V*_pmax_/*V*_cmax_ could be taken into consideration to depict a complete picture of the functional coordination (Yin *et al.*, 2016).

In C_3_ plants *J*_max_/*V*_cmax_ displays acclimation with changing environmental factors (temperature, CO_2_, light, and water availability; Onoda *et al.*, 2005; Rodriguez-Calcerrada *et al.*, 2008; Kromdijk & Long, 2016; Yin *et al.*, 2018), and it is expected that *J*_max_/*V*_cmax_ and *V*_pmax_/*V*_cmax_ might also vary in C_4_. Sage and McKown (2006) proposed that C_4_ may show less plasticity and acclimation in phenotypical traits in response to global climate change, due to their complex anatomical and biochemical features, e.g., the structural and physiological integration of the mesophyll-bundle sheath complex. Combining theoretical predictions and empirical examination of how *J*_max_/*V*_cmax_ and *V*_pmax_/*V*_cmax_ vary with environment could elucidate the acclimation capability C_4_ and further show if acclimation occurs in an optimal manner. Such an understanding of C_4_ responses to changing climate would reduce the uncertainty of global carbon flux prediction (Beerling & Quick, 1995; Quebbeman & Ramirez, 2016; Croft et al., 2017; Rogers et al., 2017).

In addition to the coordination between CB and C_4_ cycle detailed above, we expect a coordination between cyclic electron transport and total electron transport. Cyclic-electron transport, which produces ATP only, has been proposed to be enhanced C_4_ plants as compared to C_3_ plants to fulfill the extra ATP requirements of C_4_ cycle (Takabayashi *et al.*, 2005; Nakamura et al., 2013; Munekage, 2016; Munekage & Taniguchi, 2016; Yin & Struik, 2018). Chlorophyll fluorescence has been widely used for C_3_ species to estimate the *J*_max_ (Yin et al., 2009; Bellasio et al., 2016); however, chlorophyll fluorescences, representing the electron transport associated with linear electron transport, differs between C_3_ and C_4_ species. Lack of details in the proportion of linear electron transport in total electron transport in C_4_ species, therefore, hindered the use of fluorescence method to estimate electron transport in C_4_ species. We expected such a ratio will be lower in C_4_ than that in C_3_. Establishing this ratio will provide for an additional tool to estimate *J*_max_ using fluorescence measurements, as well as *V*_cmax_ and *V*_pmax_ for C_4_ species.

In the current study, we first used physiological models coupling photosynthesis, hydraulics and nitrogen stoichiometry to predict the optimal *J*_max_/*V*_cmax_ and *V*_pmax_/*V*_cmax_. Then, we performed in vivo experiments to estimate these parameters empirically on grass lineages including C_3_ and C_4_ species selected from the PACMAD clade (Grass Phylogeny Working Group II, 2012; Spriggs *et al.*, 2014) and compared these to published in vitro measurements. By sampling multiple independent origins of C_4_ species within a phylogenetic context (Edwards *et al.*, 2007; Cavender-Bares *et al.*, 2009), we were able to use phylogenetic comparative methods to examine the divergence of traits between C_3_ and C_4_ species and, to detect whether there are continuous evolutionary trends. In sum, we used optimality modeling, physiological measurements and evolutionary comparative methods to examine evolutionary trends, the approach to optimality, and to gain a better formal understanding of *J*_max_/*V*_cmax_ and *V*_pmax_/*V*_cmax_ in C_4_ photosynthesis to reduce uncertainty in modeling the carbon/nitrogen cycle, vegetation dynamics and productivity. Specifically, we aimed to test and examine the following hypotheses and questions. (1) Reorganization in resource allocation between Rubisco carboxylation and electron transport and coordination between CB cycle and C_4_ cycle in C_4_ resulted in higher *J*_max_/*V*_cmax_ and lower flr-ETR/*J*_max_ than C_3_. (2) C_4_ values for *J*_max_/*V*_cmax_ and *V*_pmax_/*V*_cmax_ are optimized the for current environment conditions. (3) Although evolving millions of years ago, selection has continued to optimize C_4_ after the evolution of the CCM, and this will yield a positive evolutionary trend between photosynthesis parameters and evolutionary age. (4) To examine the acclimation capability of C_4_ species using predictions and measurements of how *J*_max_/*V*_cmax_ and *V*_pmax_/*V*_cmax_ vary with environmental change.

## Materials and Methods

### Plant material

We cultivated 35 closely related species, including nine C_3_ and 26 C_4_. The species belong to nine independent origins of closely-related C_3_ and C_4_ lineages. Detailed cultivation information is the same as Zhou *et al.* (2020). We, then, extracted the dated phylogenetic tree for the species from the dated phylogenetic trees (Spriggs *et al.*, 2014) (Fig. S1).

### Physiological Modeling

#### Optimal *J*_max_/*V*_cmax_ and *V*_pmax_/*V*_cmax_

Based on the C_3_ and C_4_ models constructed in Zhou *et al.* (2018), which incorporates the soil-plant-air water continuum into traditional C_3_ and C_4_ photosynthesis models (Farquhar et al., 1980; von Caemmerer, 2000), we added stochiometric correlations between photosynthesis parameters and nitrogen. Using such a framework, we can model the optimal *J*_max_/*V*_cmax_ and *V*_pmax_/*V*_cmax_ simultaneously considering the following nitrogen stoichiometry:

The total nitrogen is the sum of different components (Evans, 1989):

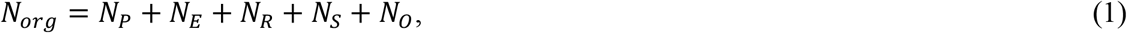

in which *N*_P_ represents the nitrogen in pigment proteins, *N*_E_ represents the nitrogen for the electron transport system, *N*_R_ represents the nitrogen of Rubisco, *N*_S_ represents nitrogen in soluble proteins except for Rubisco, and *N*_O_ represents additional organic leaf nitrogen not invested in photosynthetic functions.

In order to model the optimal *J*_max_/*V*_cmax_ and *V*_pmax_/*V*_cmax_, we need to consider the nitrogen stoichiometry among *J*_max_, *V*_cmax_ and *V*_pmax_. We used empirical relationships found in previous studies (Evans & Poorter, 2001; Niinemets & Tenhunen, 1997; Quebbeman & Ramirez, 2016):

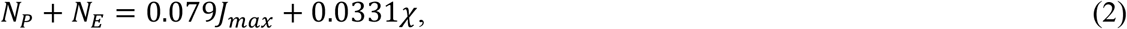

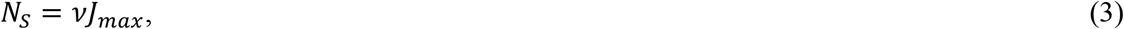

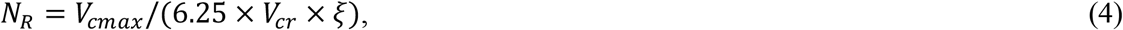

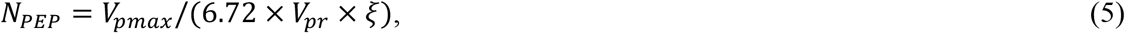

*χ* is the concentration of chlorophyll per unit area (μmol Chl m^−2^), 0.079 is in mmol N s (μmol)^−1^, and 0.0331 is in mmol N (μmol Chl)^−1^, *v* ≈ 0.3 (mmol N s (μ mol)^−1^. *V*_cr_ is the specific activity of Rubisco (the maximum rate of RuBP carboxylation per unit Rubisco; ≈ 20.5 μmol CO_2_ (g Rubisco)^−1^s^−1^) and 6.25 is grams RuBisCO per gram nitrogen in RuBisCO. *V*_pr_ is the specific activity of PEPc, that is, the maximum rate of RuBP carboxylation per unit PEPc (≈ 181.7 μmol CO_2_ (g PEPC)^−1^s^−1^), 6.72 is grams PEPc per gram nitrogen in PEPC (calculated from the amino acids composition of Fujita et al., 1984), and *ξ* is the mass in grams of one millimole of nitrogen equal to 0.014 g N (mmol N)^−1^.

Further, we simplify the equation (2) by assuming there is a coordination of resource allocation between chlorophyll and electron transport for saturated light intensity, which determines the *J*_max_. We make this assumption for the light saturated condition and use the empirical equation of Croft *et al.* (2017) to equation (2)

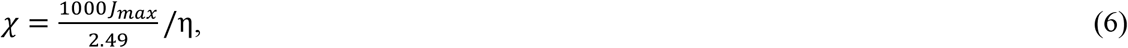

where η is the average molar mass for chlorophyll (900 g/mol). Thus,

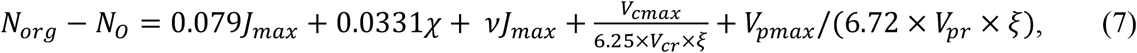

When the light intensity varies, the following function is used to adjust the electron transport rate (Ögren & Evans, 1993):

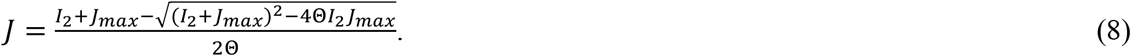

Also, all the photosynthetic parameters are temperature-sensitive (Zhou *et al.*, 2018). In the optimal modeling processes, we set *N*_org_- *N_O_* as constant of 129 mmol N m^−2^ (which yield a *V*_cmax_= 39 μmol m^−2^ s^−1^, *J*_max_= 195 μmol m^−2^ s^−1^and *V*_pmax_= 78 μmol m^−2^ s^−1^, if assuming *J*_max_/*V*_cmax_=5 and *V*_pmax_/*V*_cmax_=2). Using these models, we modeled the assimilation rates with different *J*_max_/*V*_cmax_ from 1 to 8 of 0.01 interval and different *V*_pmax_/*V*_cmax_ from 0.5 to 5 of 0.01 to find the globally optimal assimilation rate with respect to both *J*_max_/*V*_cmax_ and *V*_pmax_/*V*_cmax_. The corresponding *J*_max_/*V*_cmax_ or *J*_max_/*V*_cmax_ under the highest assimilation rates represent the optimal ratios. Then, we also model the locally optimal *J*_max_/*V*_cmax_ and *V*_pmax_/*V*_cmax_ when constraining the corresponding *V*_pmax_/*V*_cmax_ and *J*_max_/*V*_cmax_ with the average measured values respectively.

Using the model described above, we were able to model the optimal *J*_max_/*V*_cmax_ and *V*_pmax_/*V*_cmax_ under different environmental gradients: CO_2_ of 200, 300, 400, 500 and 600 ppm; VPD and ψ_S_ of (0 MPa, 0.15) (0.625, −0.5), (1.25, −1), (1.875, −1.5), and (2.5, −2); light intensity of 2000, 1600, 1200, 800 and 400 μmolm^−2^s^−1^; temperature of 15, 20, 25, 30 and 35 °C. Since there is potential uncertainty for stochiometric relationships, we performed sensitivity analysis for the total nitrogen (from 100% to 50% with 10% interval of the regular nitrogen) for optimal *J*_max_/*V*_cmax_ and *V*_pmax_/*V*_cmax_ results and we also performed sensitivity analysis for stoichiometry of PEPC by varying the 1/(6.72×*V*_pr_ × *ξ*) term in Eq. 5 from 50% to 800%. For the C_3_ pathway, all the modeling process are similar with the C_4_ and a same value of *N*_org_- *N_O_* is used, except that a simplified version of equation (7) is used as below (Quebbeman & Ramirez, 2016):

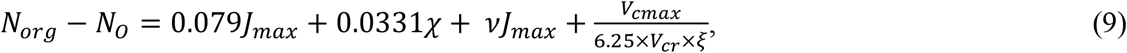

#### Effects of decreasing *V*_cmax_

Using the model, we also simulated the effect of decreasing *V*_cmax_ on the assimilation rate of both C_3_ and C_4_ pathway. In this modeling process, we hold *J*_max_ and other photosynthetic parameters constant as the initial modeling condition as above, but varying the *V*_cmax_ to 100%, 90%, 80%, 70%, 60% and 50% of the original values of C_3_.

### Gas exchange and fluorescence measurements

We measured *A/C_i_* curves using a LI-6400XT (LI-COR Inc., Lincoln, NE, USA) for all the species by setting the CO_2_ concentrations as 400, 200, 50, 75, 100, 125, 150, 175, 200, 225, 250, 275, 300, 325, 350, 400, 500, 600, 700, 800, 1000, 1200, 1400 ppm under light intensity of 2000 μmolm^−2^s^−1^. Data were recorded when the intercellular CO_2_ concentration equilibrated 2-5 minutes. Fluorescence was measured along with *A*/*C*i curves using a 2 cm^2^ fluorescence chamber head. After the change of CO_2_ concentration, the quantum yield was measured by multiphase flash when *A* reached a steady state (Bellasio *et al.*, 2014). The leaf temperatures were controlled at 25°C, VPD varied at 1-1.7kPa and the flow rate of 500 μmol s^−1^ for all the measurements. The cuvette was covered by Fun-Tak to lessen leakiness. We used the estimation method in Zhou *et al.* (2019) for *A*/*C*_i_ curves to estimate in vivo *V*_cmax_, *J*_max_, and *V*_pmax_ with one slight methodological change. Since it is thought that the region of *V*_cmax_ limit *A*/*C*_i_ is very narrow in C_4_ species, we assigned the Ci regions limited by carbonic anhydrase and *V*_cmax_ with a very low criteria of 5 Pa or below. We let the data points with *C*_i_ ranging from 5 to 60 Pa CO_2_ to be freely determined by which of the four potential limitation states to minimize the estimation error. Using this method, we avoided the potential bias of including optimal perspectives to the estimation method which could occur when directly assigning the cross points co-limited by *V*_cmax_, *V*_pmax_ and *J*_max_. Fluorescence results were used to calculate the flr-ETR (Genty *et al.*, 1989).

Furthermore, in vivo *V*_cmax_ estimated from *A/C*_i_ could be a source of uncertainty in C_4_ species. In order to provide additional support to the measured *J*_max_/*V*_cmax_ and *V*_pmax_/*V*_cmax_ value, we collected in vitro measured values for *V*_cmax_ and *V*_pmax_ from previous research, which includes eleven studies with 87 averaged results reported under current and varying environmental conditions (Supplementary Material I). Since it is impossible to obtain in vitro *J*_max_ and the estimation of *J*_max_ from *A*/*C*_i_ curves is considered reliable, we also obtain the corresponding *A*/*C*_i_ curves from these studies if they were reported, to obtain the *J*_max_. The combination of in vivo and in vitro measurements should yield a good representation of current *J*_max_/*V*_cmax_ and *V*_pmax_/*V*_cmax_ states in the C_4_ plants.

### Chlorophyll measurements and leaf nitrogen

Chlorophyll were measured using the spectrophotometer method (Porra *et al.*, 1989). We cut the fresh leaves of species into pieces of 0.5 mm long (total leaf area was measured) and submerged the fragments into DMF. After all the Chlorophyll was extracted and the leave turned white, the supernatant was used to measure the absorption under 663.8 nm and 646 nm. Total Chlorophyll concentrations were calculated using the equation of Porra *et al.* (1989). We measure leaf nitrogen contents for each sample using the CHNOS analyzer (ECS4010, Costech Analytical Technologies, Inc., Valencia, CA).

### Phylogenetic analysis

We fitted each of the photosynthetic parameters (*V*_cmax_, *J*_max_, *J*_max_/*V*_cmax_, Total Chl, flr-ETR/*J*_max_, flr-ETR, *V*_pmax_ and *V*_pmax_/*V*_cmax_) to six different evolutionary models falling into Brownian Motion model and Ornstein-Uhlenbeck Model using the R package of “mvMORPH” (Table S1). The small-sample-size corrected version of Akaike information criterion (AICc, the lower AICc, the better fit) and Akaike weights (AICw, the higher AICw, the better fit) were used as criteria to figure out the best-fitted model. We used likelihoodratio test (LRT) method to test whether one model variant performs significantly better than others and to determine whether there are significant differences between C_3_ and C_4_ species. We also extract the evolutionary ages for each C_4_ species from the dated phylogeny (Spriggs *et al.*, 2014). We regressed the above photosynthetic traits with evolutionary ages to detect potential evolutionary trends.

## Results

### In vivo and in vitro *J*_max_/*V*_cmax_ follow the global optima in C_4_, but *V*_pmax_/*V*_cmax_ does not

In vivo and in vitro *J*_max_/*V*_cmax_ are consistent with the optimal predictions under current CO_2_ conditions, but *V*_pmax_/*V*_cmax_ fell in to the optimal range under CO_2_ of 200 ppm. The global optima modelling results indicated maximal photosynthesis at the *J*_max_/*V*_cmax_ of 5-6.2, which is relatively constant across different CO_2_ concentrations, while the optimal range for *V*_pmax_/*V*_cmax_ for C_4_ species is 1.4-2.2 at CO_2_ of 200 ppm, but decreases to 1-1.6 when CO_2_ reaches 400 and 600 ppm (Fig. 1b, d, f). The averaged in vitro and in vivo *J*_max_/*V*_cmax_ are consistent with the global optimal predictions under CO_2_ of 400 ppm (Fig. 1, Fig. 2a, Supplementary Material I), as well as the locally optimal predictions controlling *V*_pmax_/*V*_cmax_ at the in vivo and in vitro level (Fig. 2). However, the averages of in vitro and in vivo *V*_pmax_/*V*_cmax_ are beyond the optimal predictions of global optima at CO_2_ of 400 ppm, but the measurement results are consistent with the optimal condition at CO_2_ of 200 ppm (Fig. 1, 3, 4, S2, Supplementary Material I). The 3D images and the contour plots also illustrate that when *J*_max_/*V*_cmax_ is at the optimal range, beyond the optimal range of *V*_pmax_/*V*_cmax_, the assimilation surface is quite flat: photosynthesis declines, but quite slowly; however, when *V*_pmax_/*V*_cmax_ drops outside of the optimal ranges, there are sharp decreases of photosynthesis (Fig. 1). In vitro measurements indicated large variations of *J*_max_/*V*_cmax_ and *V*_pmax_/*V*_cmax_ at the species level, and in vivo results, within a small scale of variation, fell completely into the range of in vitro results (Fig. 2a and 3a).

**Fig. 1.**
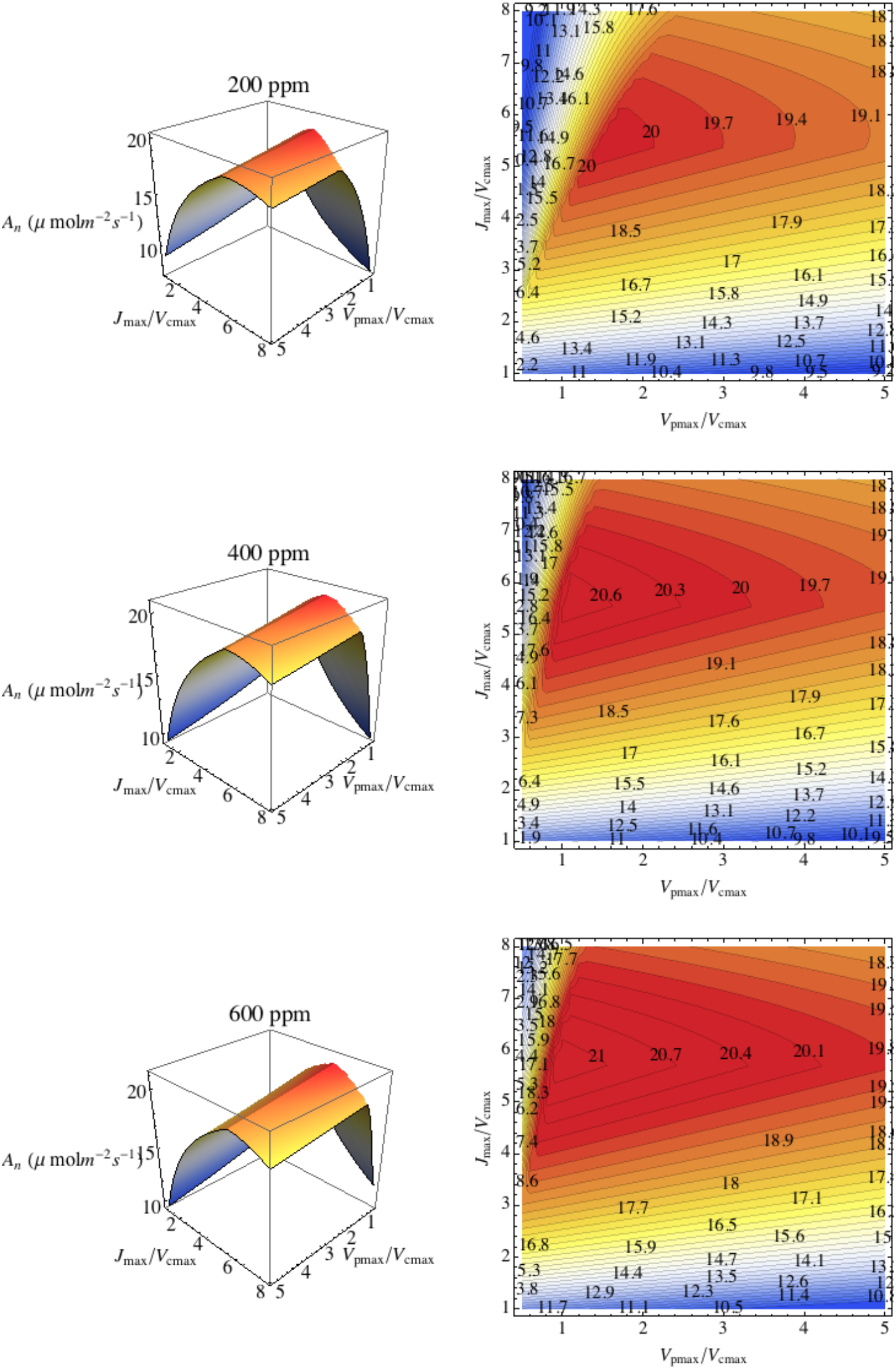
Modeling results of assimilation rate with respect to maximal electron transport to maximal Rubisco carboxylation (*J*_max_/*V*_cmax_) and maximal PEP carboxylation to maximal Rubisco carboxylation (*V*_pmax_/*V*_cmax_) under CO_2_ concentration of 200, 400 and 600 ppm. Other environmental conditions are soil water potential (ψ_S_)=-0.5 MPa, VPD=0.625, temperature of 25 °C and light intensity of 2000 μmol m^−2^ s^−1^. Left: 3D plot; right: corresponding contour plot.

**Fig. 2.**
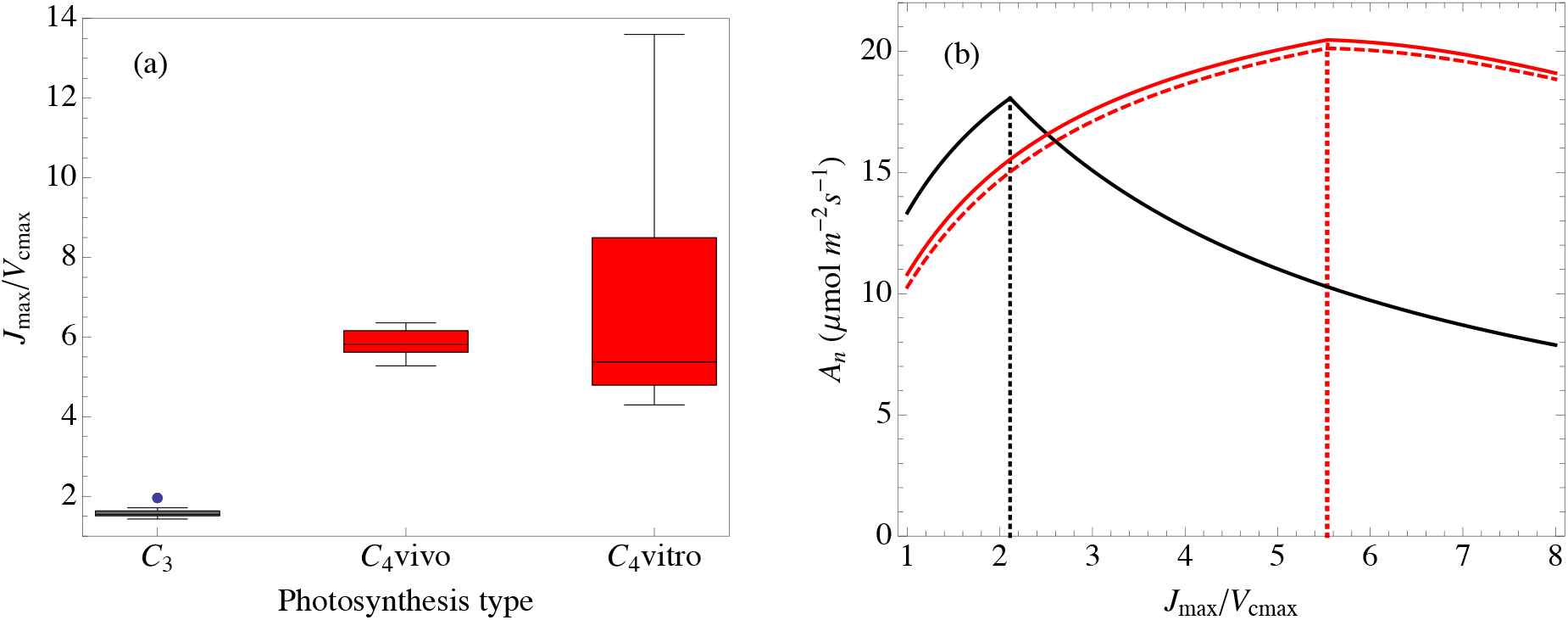
Empirical measurements (a) and optimal modeling results (b) of *J*_max_/*V*_cmax_ for C_3_ and C_4_ under ψ_S_=-0.5 MPa, VPD=0.625, temperature of 25 °C and light intensity of 2000 μmol m^−2^ s^−1^, the cultivating environmental condition. In (b), the black line represents C_3_, solid red line represents C_4_ modeling results with controlling *V*_pmax_/*V*_cmax_ at the in vivo measurement level, dashed red line represents C_4_ modeling results with controlling *V*_pmax_/*V*_cmax_ at the in vitro measurement level.

**Fig. 3.**
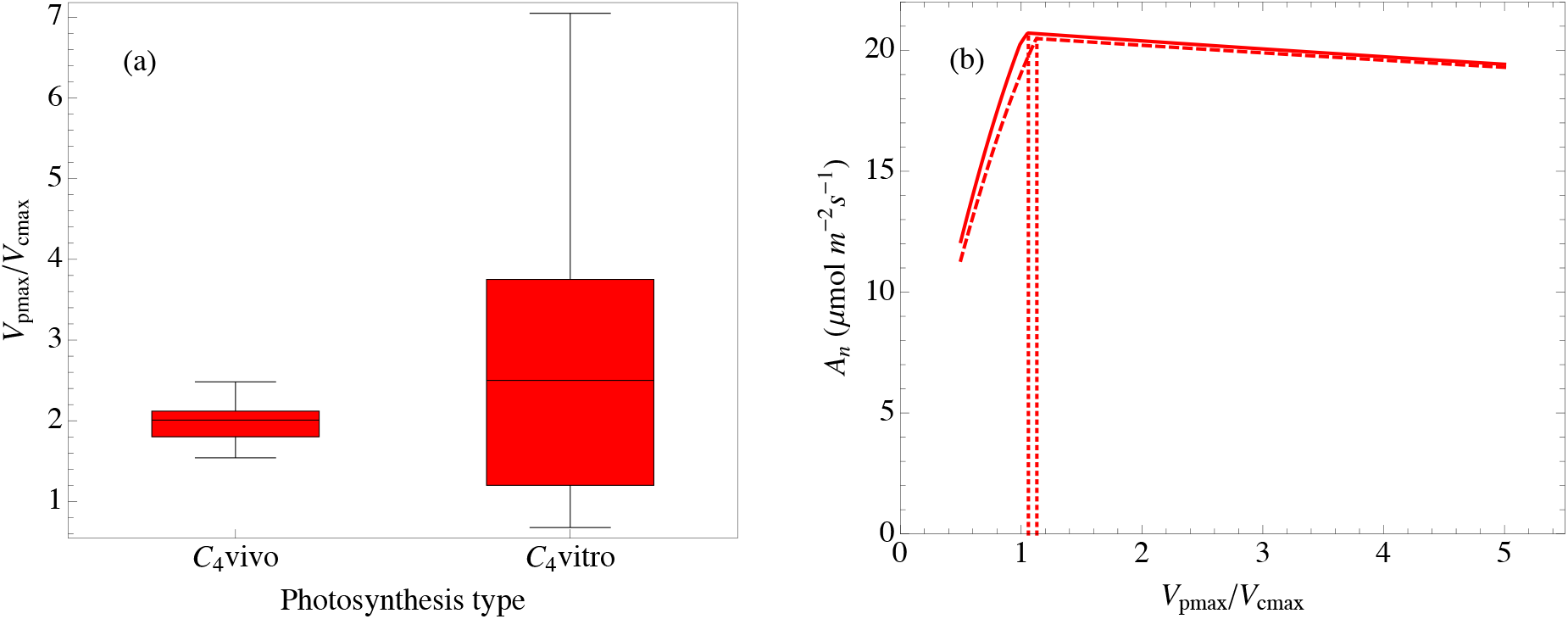
Empirical measurements (a) and optimal modeling results (b) of *V*_pmax_/*V*_cmax_ for C_4_ under ψ_S_=-0.5 MPa, VPD=0.625, temperature of 25 °C and light intensity of 2000 μmol m^−2^ s^−1^, the cultivating environmental condition. In (b), solid red line represents C_4_ modeling results with controlling *J*_max_/*V*_cmax_ at the in vivo measurement level, dashed red line represents C_4_ modeling results with controlling *J*_max_/*V*_cmax_ at the in vitro measurement level.

**Fig. 4.**
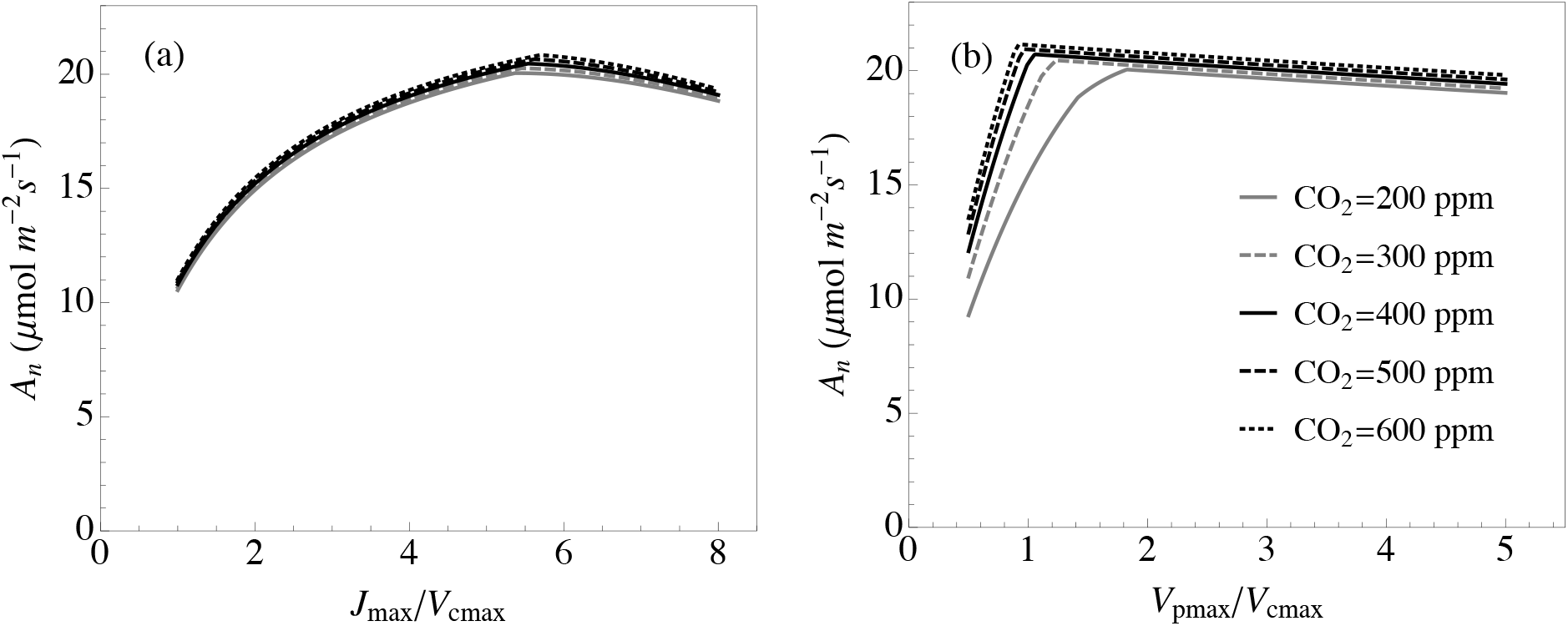
Modeling results of assimilation rate with varying *J*_max_/*V*_cmax_ and *V*_pmax_/*V*_cmax_ for C_4_ under different CO_2_ concentration and ψ_S_=-1MPa, VPD=1.25, temperature of 25 °C and light intensity of 2000 μmol m^−2^ s^−1^, the common grassland growth condition. Modeling results were obtained by controlling the other parameter at the in vivo measurement level.

### C_4_ species have higher J_max_/V_cmax_ and lower *flr-ETR*/J_max_ than C_3_ species

Phylogenetic analysis shows the *J*_max_/*V*_cmax_ follows the Ornstein-Uhlenbeck model with a higher stable state for C_4_ species and a lower stable state for C_3_ (Table 1; Fig. 1a). Such an empirical relationship is consistent with the optimal modeling predictions for C_3_ and C_4_. We looked further into how such a higher *J*_max_/*V*_cmax_ in C_4_ species is reached by comparing individual empirical parameters. C_4_ species has significantly higher stable states of *J*_max_, but significantly lower stable states of *V*_cmax_ and nitrogen content than their closely-related C_3_ (Table 1). In order to examine the potential effects of decreasing *V*_cmax_ on assimilation rate, we held the *J*_max_ as constant and changed the *V*_cmax_ from 100% to 50% of the original C_3_ parameter values in the C_3_ and C_4_ models. A decrease in *V*_cmax_ will significantly decrease the assimilation rates of C_3_ species from 10 °C to 35°C under different CO_2_ concentrations, while decreasing *V*_cmax_ has little effects on the assimilation rates of C_4_ species (Fig. 5).

**Fig. 5.**
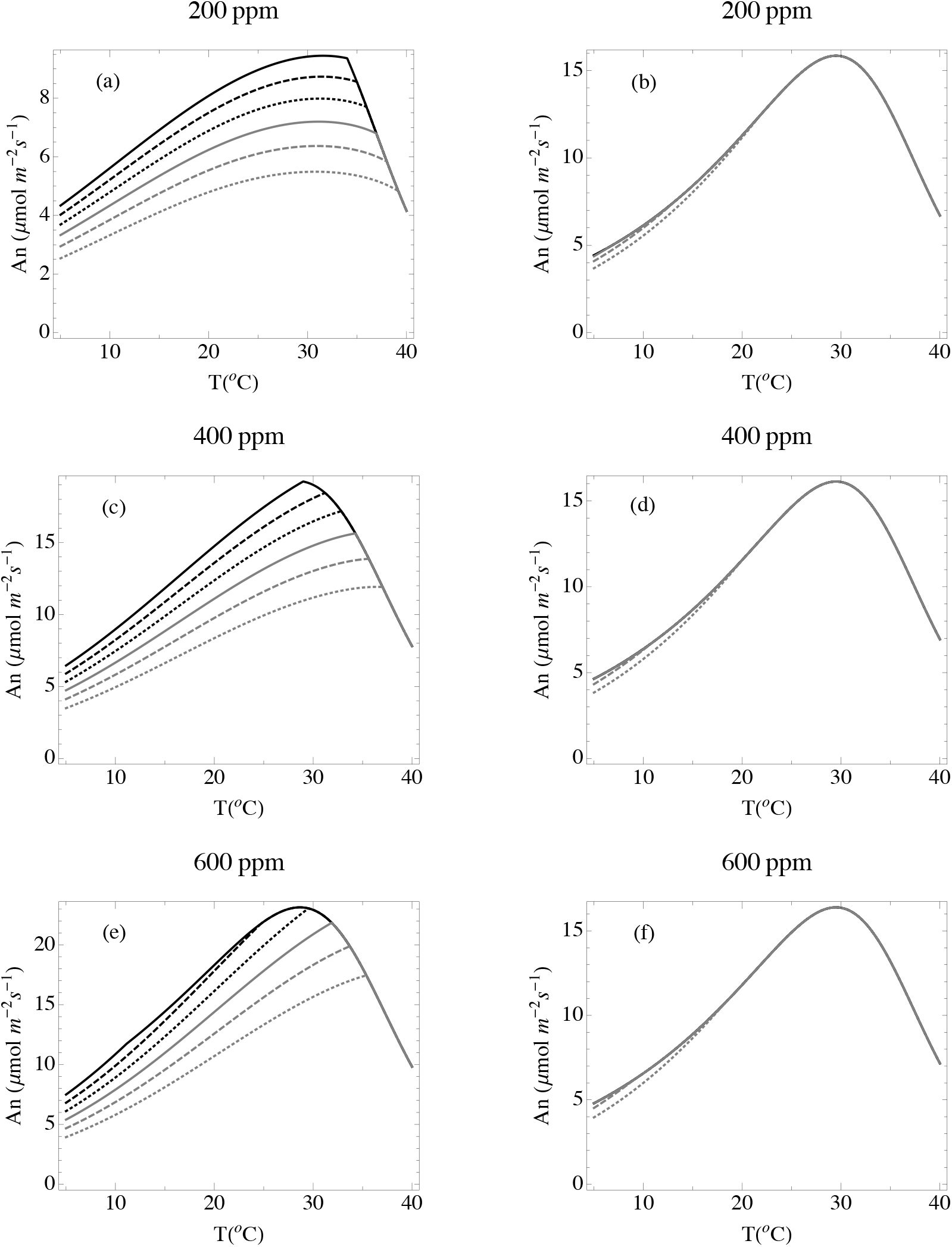
Modeling results of changing *V*_cmax_ on assimilation rates for C_3_ (acd) and for C_4_ (bdf) under different CO_2_. Solid black line: the initial modeling condition of *V*_cmax_ (a typical C_3_ value of 69 μmol m^−2^ s^−1^); dashed black line: 90% of the initial *V*_cmax_; dotted black line: 80% of the initial *V*_cmax_; solid grey line: 70% of the initial *V*_cmax_; dashed grey line: 60% of the initial *V*_cmax_; dotted grey line: 50% of the initial *V*_cmax_.C_3_ and C_4_ parameters shared similar parameters except for the carbon concentration mechanism and different hydraulic conductance to mimic the initial origin of C_4_ (see Zhou et al (2020) for other parameters).

**Table 1.**
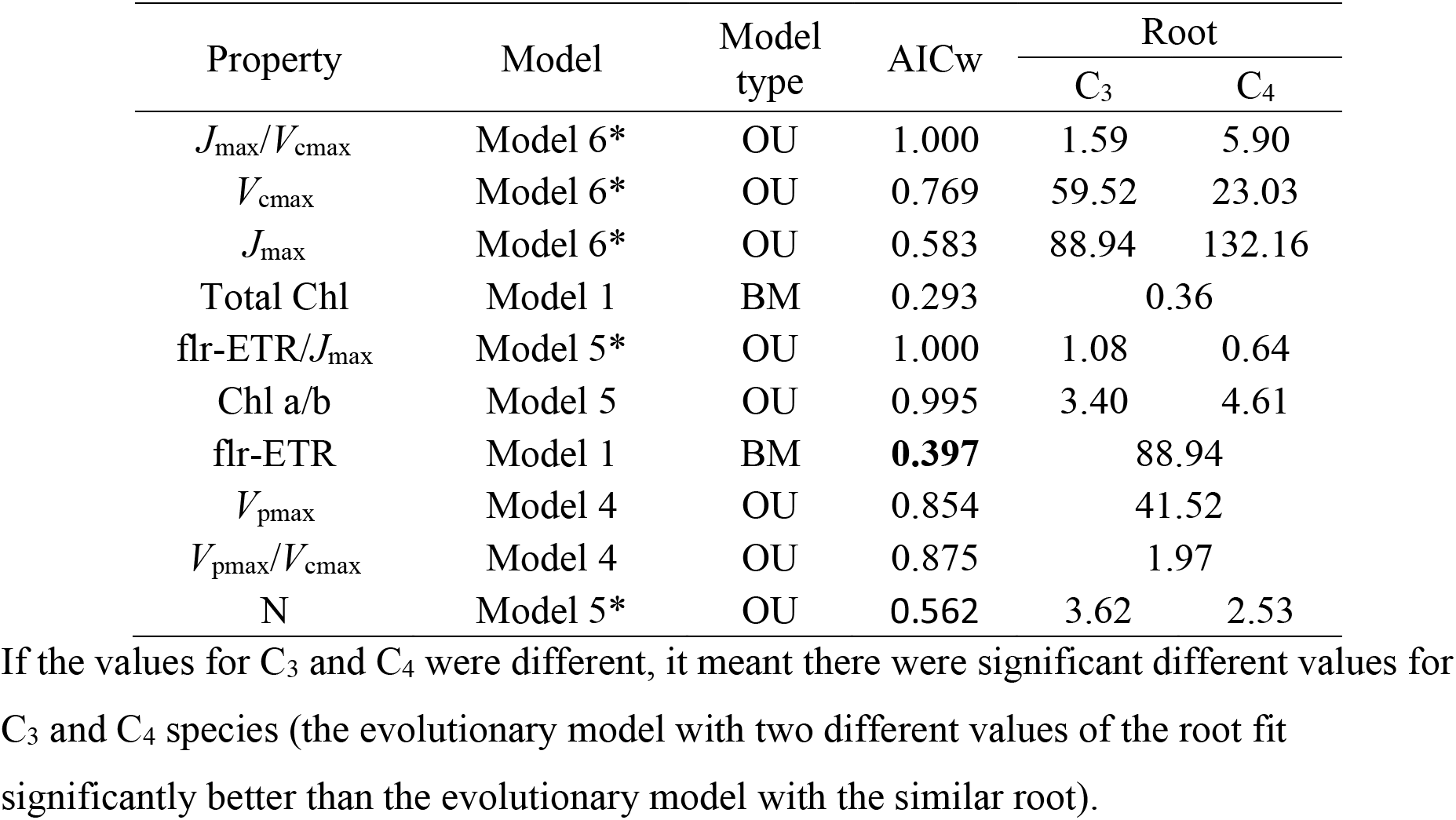
Phylogenetic results of the best-fitted models and their parameters for photosynthesis parameters (summarizing Table S2-S10).

The flr-ETR/ *J*_max_ was significantly higher in C_3_ species than that in their closely related C_4_ species, with the stable states of 1.08 and 0.64 for C_3_ and C_4_ respectively (Table 1). Phylogenetic analysis indicated C_3_ and C_4_ species had the similar stable states of ETR, but C_4_ has a higher total electron transport rate, *J*_max_ (Table 1).

### The positive evolutionary trends of photosynthetic parameters

Plotting the photosynthetic parameters with evolutionary ages, extracted from a dated phylogeny for the multiple lineages, allows us to look for further evolutionary trends in C_4_ and their closely related C_3_ species. Regressions of evolutionary age versus photosynthetic traits provide signals for long-term directional trends in photosynthetic machinery following the establishment of C_4_ photosynthesis (Fig. 6). *A*_max_, *J*_max_, total chlorophyll, *V*_cmax_ and *V*_pmax_ showed significant positive correlations with evolutionary age in C_4_ species, but not C_3_ species, while nitrogen, *J*_max_/*V*_cmax_, *V*_pmax_/*V*_cmax_ and flr-ETR/*J*_max_ did not show significant correlation with the evolutionary age.

**Fig. 6.**
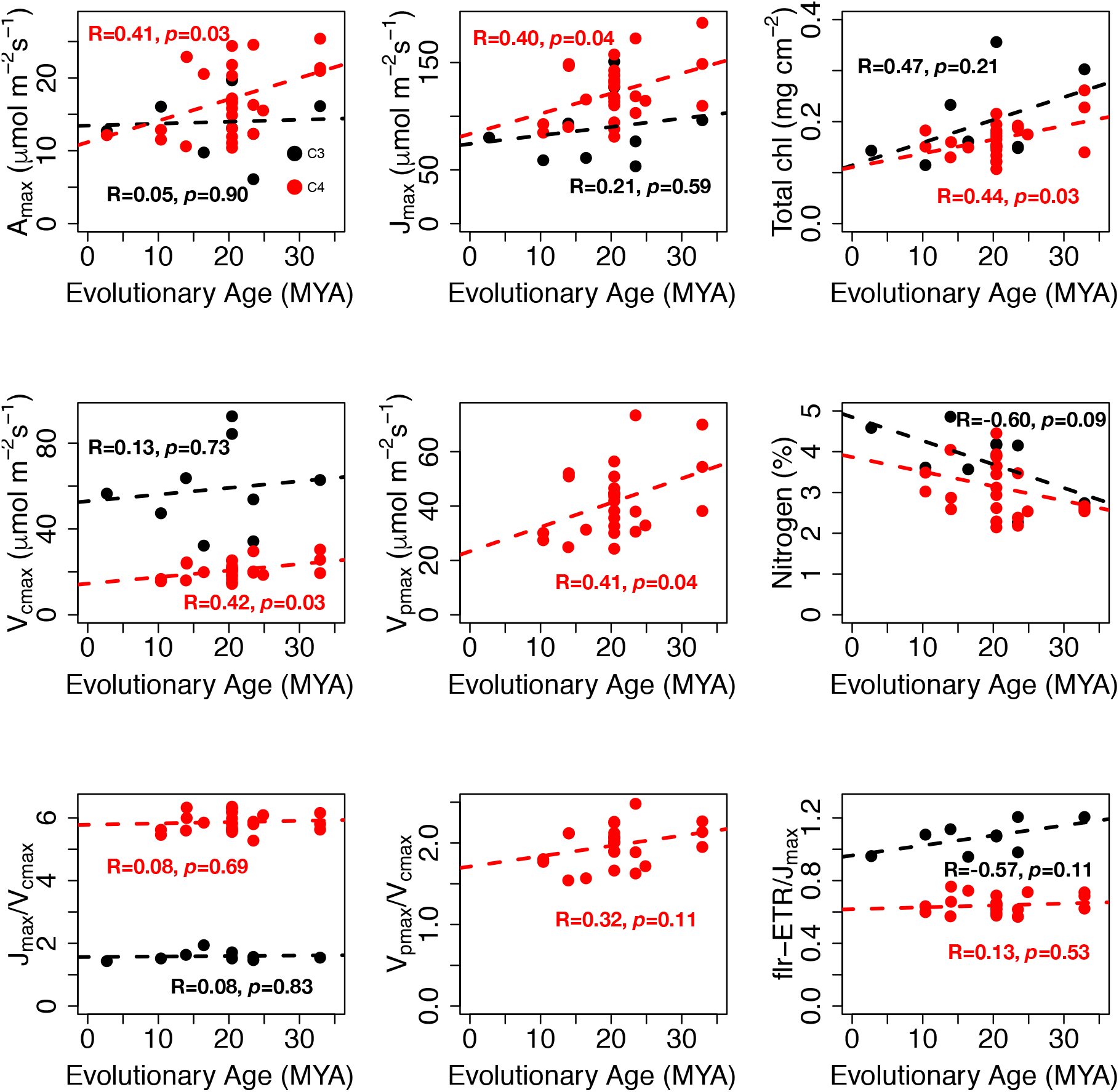
The regression for maximal assimilation rate (*A*_max_), *J*_max_, total cholorophyll (Total chl), *V*_cmax_, *V*_pmax_, nitrogen concentration, *J*_max_/*V*_cmax_, *V*_pmax_/*V*_cmax_ and fluorescence-estimated electron transport to *J*_max_ (ETR/ *J*_max_) vs. the evolutionary age for the nine origins to show the evolutionary trend within C_4_ (red) and within their closely-related C_3_ species (black).

### Optimal variation of J_max_/V_cmax_ *and* V_pmax_/V_cmax_ with environmental conditions

To understand how the *J*_max_/*V*_cmax_ and *V*_pmax_/*V*_cmax_ varied theoretically in response to environmental changes (Fig. 7a, b, Fig. S3a, b), we calculated their optimal value for varying CO_2_ concentrations, water limitations, temperatures, and light intensities. The optimal *J*_max_/*V*_cmax_ is predicted to increase linearly in C_3_ and, but increase a little bit and maintain relatively constant in C_4_ species with increasing CO_2_ concentration (Fig. 7a). The optimal *J*_max_/*V*_cmax_ decreases similarly in both C_3_ and C_4_ species along with increasing water limitation and increases similarly with decreasing light intensity and increasing temperatures (Fig. 7). The changes of *J*_max_/*V*_cmax_ with water limitation, light intensity, and temperature are non-linear, with the rate-of-change increasing greatly after a threshold (water limitation of ψ_S_=−1, VPD=1.25, the light intensity of 800 μmol m^−2^ s^−1^ and temperature of 30 °C). The optimal *V*_pmax_/*V*_cmax_ decreases along with the increase of the CO_2_ concentration, especially when CO_2_ increases from 200 ppm to 300 ppm, but the change is little when CO_2_ is above 400 ppm (Fig. 7c, S3c). However, *V*_pmax_/*V*_cmax_ is relatively constant with the varying of water limitation conditions and light intensity (Fig. 7c,d, S3c,d). *V*_pmax_/*V*_cmax_ decreases with the rise in temperature from 15 to 35 °C (Fig. 7d, S3d).

**Fig. 7.**
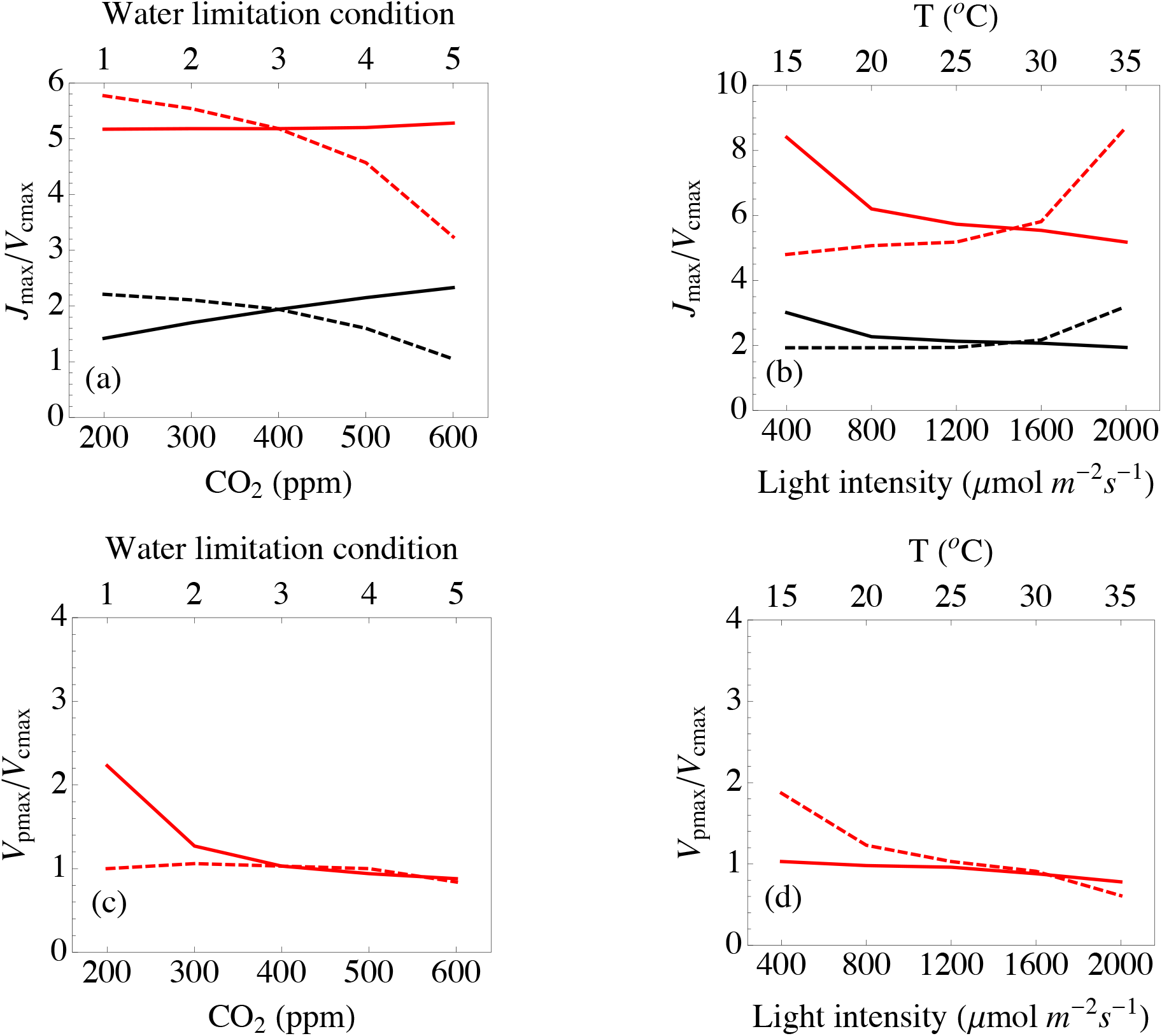
Modeling results of optimal *J*_max_/*V*_cmax_ and *V*_pmax_/*V*_cmax_ for C_3_ (black lines) and C_4_ (red lines) under different environmental conditions. (a) Solid line: different CO_2_; dashed line: different water limitation conditions (1: saturated water; 2: ψ_S_=-0.5 MPa, VPD=0.625; 3: ψ_S_=-1 MPa, VPD=1.25 MPa; 4: ψ_S_=-1.5 MPa, VPD=1.875; 5: ψ_S_=-2 MPa, VPD=2.5). (b) Solid line: different light intensities; dashed line: different temperature. Modeling results were obtained by controlling the other parameter at the in vivo measurement level.

### Sensitivity analysis for optimal *J*_max_/*V*_cmax_ and *V*_pmax_/*V*_cmax_

Since there is a great variation of total nitrogen content in plants and there is uncertainty in the stoichiometry of PEPC, we performed sensitivity analysis and found the optimal modeling of *J*_max_/*V*_cmax_ and *V*_pmax_/*V*_cmax_ are robust. For C_3_ photosynthesis, optimal *J*_max_/*V*_cmax_ increases slightly with decreasing total nitrogen (7.7% increase of *J*_max_/*V*_cmax_ with 50% reduction in total nitrogen), while for C_4_, optimal *J*_max_/*V*_cmax_ decreases with total nitrogen (16.2% increase of *J*_max_/*V*_cmax_ with 50% reduction in total nitrogen) (Fig. S4a). The optimal *V*_pmax_/*V*_cmax_ increases somewhat with decreasing total nitrogen (8.7% increase of *V*_pmax_/*V*_cmax_ with 50% reduction in total nitrogen) (Fig. S4a). The optimal *J*_max_/*V*_cmax_ is relatively constant with the change of stoichiometry of PEPC from 50% to 200%, but increases as the stoichiometry increases from 200% to 800% (Fig. S3b). The optimal *V*_pmax_/*V*_cmax_ is relatively robust with the change of stoichiometry of PEPC.

## Discussion

Our modeling efforts provide an explanation for the observed variation in *J*_max_/*V*_cmax_ and *V*_pmax_/*V*_cmax_, and why *V*_pmax_/*V*_cmax_ appears to be optimized for the lower bounds of atmospheric CO_2_ of the Pleistocene. Our reported value of *V*_pmax_/*V*_cmax_ are comparable with previous studies (Kubien *et al.*, 2003; Pengelly *et al.*, 2010; Yin *et al.*,2011, 2016; Pignon and Long, 2020), and two recent papers also indicated that the coordination between CB and C_4_ cycles might represent a legacy of ancient low CO_2_ conditions (Sundermann et al., 2018; Pignon and Long, 2020). All extant C_4_ species have gone through a low CO_2_ bottleneck over the last 5 million years (Edwards et al., 2010). This bottleneck may have resulted in a strong selection to increase *V*_pmax_/*V*_cmax_ to maintain a high assimilation rate under the low CO_2_ of glacial maxima (~200 ppm). As CO_2_ has risen, first with the beginning of the Holocene interglacial, and then again with the continual burning of fossil fuels, *V*_pmax_/*V*_cmax_ remained constant and consequently exceeded the optimal *V*_pmax_/*V*_cmax_ at higher CO_2_. The effects of a too-high *V*_pmax_/*V*_cmax_ on assimilation rate are, however, minimal, thus the selection against increasing *V*_pmax_/*V*_cmax_ was likely weak. The explanation directly rests on the topology of assimilation surface: when *J*_max_/*V*_cmax_ and *V*_pmax_/*V*_cmax_ are lower than the optimal states, assimilation rate declines greatly; but when *J*_max_/*V*_cmax_ and *V*_pmax_/*V*_cmax_ exceed the optimal states, the decrease of assimilation rate is minimal. Similarly, if a high *J*_max_/*V*_cmax_ was selected for a given species under, for example, growth at relatively low light conditions, the high *J*_max_/*V*_cmax_ could well be maintained for a long time even when moving to a high light environment. Such small changes in assimilation rate may not change fitness enough to drive evolution toward optimal states in natural plants; artificial selection and manipulation to change the *J*_max_/*V*_cmax_ and *V*_pmax_/*V*_cmax_ toward the optimal states, however, might show potential in regard to increasing total assimilation rate and productivity (Walker et al., 2018; Pignon and Long, 2020).

It has been hypothesized that after the evolution of the full C_4_ CCM that selection across various habitats would select for further physiologically optimal adjustments (Williams et al., 2013; Stata et al., 2019); we find unequal support for such adjustments through time as several physiological measures were positively related to evolutionary age, but there were no trends with photosynthetic coordination. From this, we conclude that initially there was very strong selection for coordination between the CB cycle, light reactions and the C_4_ CCM, and that changes in secondary or tertiary traits led to an increase in, for example, maximum CO_2_ assimilation rate through time. Zhou et al. (2020) illustrated that hydraulic traits could be just such an example of the secondary traits: the formation of C_4_ pathway allowed for lower stomatal conductance which triggered the decline of leaf hydraulic conductance and capacitance over evolutionary time to further maximize the assimilation rate. Additionally, anatomical organization within the leaf, like the ratio of mesophyll cells to bundle-sheath cells, the 3D arrangement of cells and shifts in intercellular airspace could also be selected upon through time towards a more optimal C_4_ photosynthetic machine (Wang et al., 2017; Alonso-Cantabrana et al., 2018; Edwards, 2019).

Higher *J*_max_/*V*_cmax_ in C_4_ than that in C_3_ indicated a change in resource allocation, namely nitrogen, between the light reactions and the CB cycle, and as a crucial evolutionary step for elevating C_4_ efficiency, it is important to look at how the reallocation may have occurred and the larger-scale consequences. The modeling results indicate that a decrease of Rubisco content is favored in C_4_, because overall nitrogen requirements decrease and such a reduction has minimal effects on net assimilation rate. Significantly lower *V*_cmax_ in all of our C_4_ species and lower Rubisco in previous studies confirmed the assertion (Brown, 1978; Ku *et al.*, 1979; Sage and Pearcy, 1987; Sharwood *et al.*, 2016). Any surplus nitrogen not invested in Rubisco could be distributed among three broad categories: 1) reallocated to the light reactions or 2) stored or used to construct new tissues or 3) simply not taken up from the growth environment, thus reducing total plant nitrogen requirements. Tissue *et al.* (1995) and Ghannoum *et al.* (2010) detected lower Rubisco content and higher chlorophyll and thylakoid content, supporting resource reallocation from RuBP carboxylation to electron transport within the leaf. Our measurements provide evidence that the coordination of *J*_max_/*V*_cmax_ resulted from a mix of hypotheses 1) and 3). The significantly higher *J*_max_ and lower *V*_cmax_ in C_4_ than their closely-related C_3_ species supports a reallocation of hypothesis 1). In addition, hypothesis 3) likely occurred together with hypothesis 1) because C_4_ species have significantly lower nitrogen content. Hypothesis 2), not exclusive to hypotheses 1) and 3), could be supported by evidence that C_4_ plants maintain larger leaf areas (Ripley *et al.*, 2008). These hypotheses are connected to potential ecological ramifications. Firstly, in a nitrogen-depleted habitat, C_4_ could have a competitive advantage as confirmed by Ripley *et al.* (2008; although Sage & Pearcy, 1987, found no evidence for this). In habitats where nitrogen is not limiting, the excess nitrogen could be used to construct more leaf area (Sage & Pearcy, 1987; Anten *et al.*, 1995; Ripley *et al.*, 2008), and greater leaf area in the early stages of growth was indeed seen by Atkinson *et al.* (2016). On the other hand, the lack of nitrogen reallocation from the CB cycle to the light reactions may indicate physiological constraints in fertile habitats. For example, photorespiration in C_3_ plants is proposed to enhance the nitrate metabolism (Oaks, 1994; Rachmilevitch *et al.*, 2004; Bauwe *et al.*, 2010; Bloom, 2015), therefore the formation of CCM, which inhibits photorespiration, may reduce overall plant-available nitrogen for C_4_. In addition, the increase of *J*_max_ in C_4_ is due to an enhanced cyclic electron transport, while maintaining the linear electron transport at the same level of C_3_. Elevating cyclic electron transport is therefore a potentially important step in engineering C_4_ photosynthesis into C_3_ crops.

Empirical evidence indicates adaptive plasticity (or acclimation) of C_4_ photosynthetic coordination occurs under different environmental conditions, consistent with optimal predictions. Contrary to what Sage and McKown (2006) proposed, C_4_ exhibited significant acclimation capability with varying CO_2_ (Pinto et al. 2014; Pinto et al. 2016), water availability (Sharwood et al. 2014), light intensity (Sharwood et al. 2014; Pengelly et al. 2010; Sonawane, 2016) and temperature (Sonawane, 2016; Pittermann and Sage 2001; Kubien and Sage 2004; Serrano-Romero and Cousins, 2020) in both *J*_max_/*V*_cmax_ and *V*_pmax_/*V*_cmax_ (Supplementary Material I). The varying trends in *J*_max_/*V*_cmax_ are generally consistent between the empirical measurements and optimal modeling predictions in the current study: higher resource allocation to electron transport under shade, elevated CO_2_, high water availability and higher temperature. However, C_3_ and C_4_ species might respond differently to environmental variations, especially in response to CO_2_ variation shown in the optimal modeling results. The *V*_pmax_/*V*_cmax_ exceededs the optimal prediction when acclimated to the current environmental conditiona, perhaps because *V*_pmax_/*V*_cmax_ is constraint by a legacy of acclimation to historical CO_2_. These acclimation responses should be considered and included in ESMs and LSMs to predict responses of C_3_ and C_4_ ecosystems to future climate change (Rogers et al., 2017; Smith and Keenan, 2020).

Our improved estimates of *J*_max_/*V*_cmax_, *V*_pmax_/*V*_cmax_ and flr-ETR/*J*_max_ for C_4_ plants could directly benefit terrestrial biosphere models. Although *J*_max_, *V*_cmax_ and *V*_pmax_ are key input parameters in global-scale models (Beerling & Quick, 1995; Zaehle *et al.*, 2005; Bonan *et al.*, 2011; Walker *et al.*, 2014), it is difficult and perhaps not feasible to measure all parameters for numerous sites. If one the ratioed parameters described here, either *J*_max_/*V*_cmax_, *V*_pmax_/*V*_cmax_ and/or flr-ETR/*J*_max_, is obtained then other parameters could be estimated. Using *J*_max_/*V*_cmax_, *V*_pmax_/*V*_cmax_ and flr-ETR/*J*_max_ is especially crucial in C_4_ species because in vivo estimation of *V*_cmax_ and *V*_pmax_ are more difficult and less reliable, and in vitro measurements are not easily performed over broad taxonomic or spatial scales. Our results suggest a value of around 5.5 for *J*_max_/*V*_cmax_ and a value of 2 for *V*_pmax_/*V*_cmax_ supported by optimality modeling and empirical measurements. With an empirical value of around 60% found in the current study, the measurement of flr-ETR could provide another tool to further estimate *J*_max_, *V*_cmax_ and *V*_pmax_. Previous theoretical analysis also inferred values of flr-ETR/*J*_max_ of 60% and 50% in C_4_ (Yin et al., 2011; Yin & Struik, 2012; Yin & Struik, 2018).

The evolution of C_4_ photosynthesis required the reorganization and coordination of the CB-cycle, the light reactions and the PEPc-based CCM. Strong divergence in *J*_max_/*V*_cmax_ between C_4_ and C_3_ species indicates the resource allocation changes between light reactions and the CB cycle that were necessary to support the enhanced ATP requirement of the C_4_ CCM (Osborn & Sack, 2012; Zhou et al., 2018). Observed *J*_max_/*V*_cmax_ were within the predicted optimal zone suggesting that the resource reallocation between Rubisco carboxylation and electron transport are operating near optimality under current environmental conditions; however, the long tail exceeding the optimal *J*_max_/*V*_cmax_ in empirical measurements indicates multiple species have over-allocated to electron transport, perhaps a legacy of native ecological conditions. The coordination between CB and C_4_ cycles was in line with the optimal conditions under 200 ppm representing an over-allocation of resources for current environmental conditions, but there is little costs to assimilation rate due to this lack of optimality. Rapid coordination occurred early in C_4_ evolution, but it appears that C_4_ photosynthesis is still under selection for further optimization. The enhanced understanding of the evolution-based photosynthetic reorganization and coordination in C_4_ photosynthesis, along with our ratio-based approach to obtain photosynthetic parameters can lead to a better parameterization of terrestrial biosphere models for C_4_.

## Acknowledgements

We sincerely thank Dr. Matteo Detto (Princeton University) for his help in building the optimal models for *J*_max_/*V*_cmax_.

