## Supplementary material for "Optimal coordination and reorganization of photosynthetic properties in C_4_ grasses": Fig. S1, Fig. S2, Fig. S3, Fig. S4

**Article title: Optimal coordination and reorganization of photosynthetic properties in $C_4$ grasses**

**Fig. S1** Phylogenetic sampling of the species for measuring photosynthetic traits and the independent evolutionary lineages corresponding to grass lineages. Black represents $C_3$ species and red represents $C_4$ species.

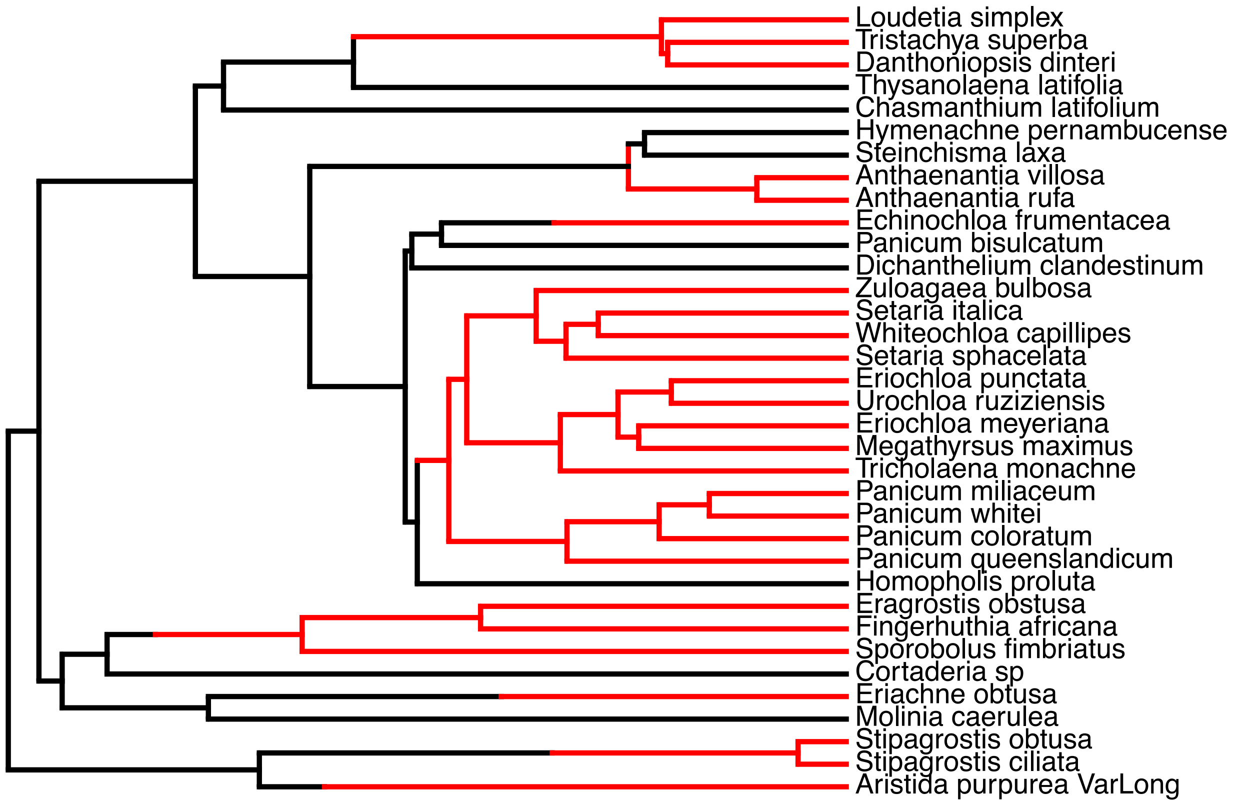

**Fig. S2** Modeling results of assimilation rate with varying $J_{max}/V_{cmax}$ and $V_{pmax}/V_{cmax}$ for $C_4$ under different $CO_2$ concentration and $\psi_S$=-1MPa, VPD=1.25, temperature of 25 °C and light intensity of 2000 μmol $m^{-2}$ $s^{-1}$, the common grassland growth condition. Modeling results were obtained by controlling the other parameter at the in vitro measurement level.

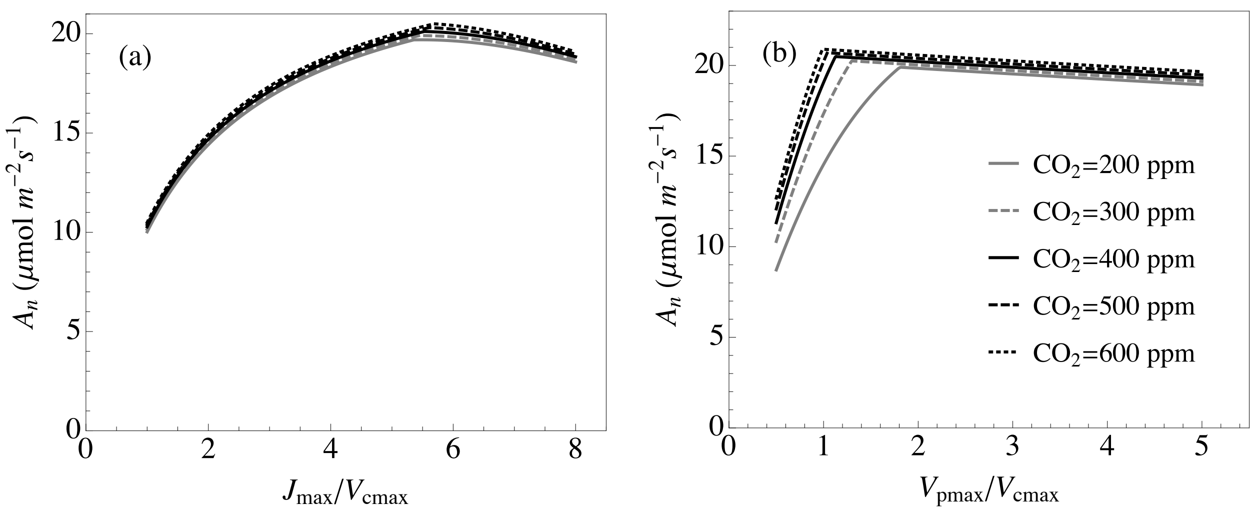

**Fig. S3** Modeling results of optimal $J_{max}/V_{cmax}$ and $V_{pmax}/V_{cmax}$ for $C_3$ (black lines) and $C_4$ (red lines) under different environmental conditions. (a) Solid line: different $CO_2$; dashed line: different water limitation conditions (1: saturated water; 2: $\psi_S$=-0.5 MPa, VPD=0.625; 3: $\psi_S$=-1 MPa, VPD=1.25 MPa; 4: $\psi_S$=-1.5 MPa, VPD=1.875; 5: $\psi_S$=-2 MPa, VPD=2.5). (b) Solid line: different light intensities; dashed line: different temperature. Modeling results were obtained by controlling the other parameter at the in vitro measurement level.

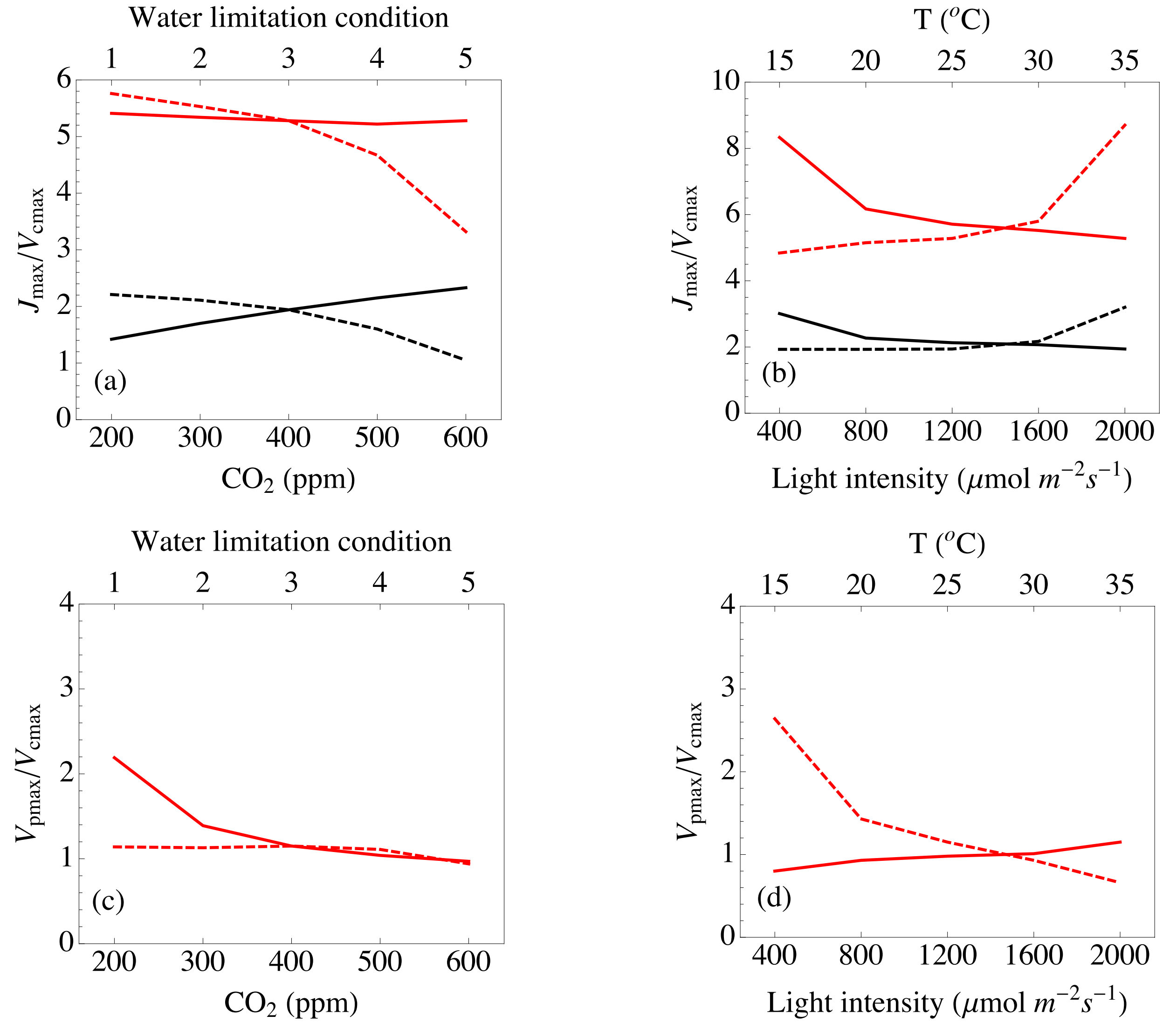

**Fig. S4** Sensitivity analysis of total nitrogen and PEPC stoichiometry on optimal $J_{max}/V_{cmax}$ for $C_3$ (black dots) and $J_{max}/V_{cmax}$ (red dots) and $V_{pmax}/V_{cmax}$ (red diamonds) for $C_4$.

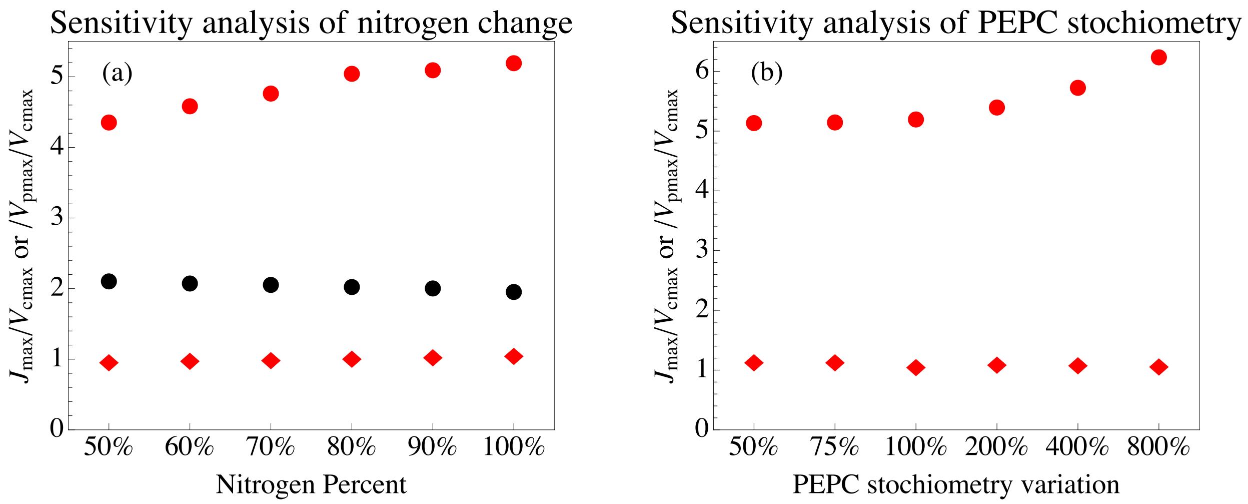

**Table S1** The evolutionary models used for the phylogenetic analysis for photosynthetic properties evolutionary differences between C3 and C4 species.

| Model | Evolution | Fluctuation rate | Root |
|---|---|---|---|
| Model 1 | BM | One | One |
| Model 2 | BM | Two | One |
| Model 3 | BM | One | Two |
| Model 4 | BM | Two | Two |
| Model 5 | OU | One | One |
| Model 6 | OU | One | Two |

**Table S2** Phylogenetic analysis results for Jmax/Vcmax for C3 and C4 species.

| Model | AICw | AIC | AICc | Variance | | Mean | |
|---|---|---|---|---|---|---|---|
| | | | | $C_3$ | $C_4$ | $C_3$ | $C_4$ |
| Model 1 | 0.000 | 138.66 | 139.04 | 0.127 | | 4.349 | |
| Model 2 | 0.000 | 103.43 | 104.20 | 0.445 | .003 | 5.520 | |
| Model 3 | 0.000 | 140.01 | 140.78 | 0.124 | | 4.028 | 4.469 |
| Model 4 | 0.000 | 104.15 | 105.49 | 0.440 | 0.002 | 5.395 | 5.557 |
| Model 5 | 0.000 | 142.80 | 143.57 | 0.188 | | 4.375 | |
| **Model 6** | **1.000** | **16.41** | **17.75** | **0.0276** | | **1.593** | **5.899** |

**Table S3** Phylogenetic analysis results for Vcmax for C3 and C4 species.

| Model | AICw | AIC | AICc | Variance | | Mean | |
|---|---|---|---|---|---|---|---|
| | | | | $C_3$ | $C_4$ | $C_3$ | $C_4$ |
| Model 1 | 0.000 | 306.21 | 306.58 | 14.801 | | 32.133 | |
| Model 2 | 0.167 | 273.40 | 274.17 | 49.056 | 0.916 | 24.216 | |
| Model 3 | 0.000 | 308.60 | 308.37 | 14.536 | | 35.451 | 30.893 |
| Model 4 | 0.064 | 275.31 | 276.65 | 49.188 | 0.908 | 23.826 | 24.345 |
| Model 5 | 0.000 | 310.22 | 311.00 | 32.372 | | 32.725 | |
| **Model 6** | **0.769** | **270.34** | **271.67** | **127.883** | | **59.524** | **20.782** |

**Table S4** Phylogenetic analysis results for Jmax for C3 and C4 species.

| Model | AICw | AIC | AICc | Variance | | Mean | |
|---|---|---|---|---|---|---|---|
| | | | | $C_3$ | $C_4$ | $C_3$ | $C_4$ |
| Model 1 | 0.201 | 338.96 | 339.33 | 33.249 | | 110.379 | |

| | | | | | | | |
|---|---|---|---|---|---|---|---|
| Model 2 | 0.074 | 340.95 | 341.73 | 33.985 | 32.687 | 110.690 | |
| Model 3 | 0.079 | 340.82 | 341.59 | 33.145 | | 107.932 | 111.346 |
| Model 4 | 0.029 | 342.81 | 344.15 | 34.153 | 32.377 | 108.327 | 111.775 |
| Model 5 | 0.032 | 342.61 | 343.38 | 66.102 | | 111.266 | |
| **Model 6** | **0.583** | **336.83** | **338.16** | **83.786** | | **88.935** | **132.164** |

**Table S5** Phylogenetic analysis results for total chlorophyll for C3 and C4 species.

| Model | AICw | AIC | AICc | Variance | | Mean | |
|---|---|---|---|---|---|---|---|
| | | | | $C_3$ | $C_4$ | $C_3$ | $C_4$ |
| **Model 1** | **0.293** | **-58.9** | **-58.6** | **0.0003** | | **0.364** | |
| Model 2 | 0.251 | -58.6 | -57.8 | 0.0006 | 0.0003 | 0.366 | |
| Model 3 | 0.285 | -58.9 | -58.1 | 0.0003 | | 0.395 | 0.352 |
| Model 4 | 0.152 | -57.6 | -56.3 | 0.0005 | 0.0003 | 0.389 | 0.357 |
| Model 5 | 0.011 | -52.3 | -51.5 | 0.0012 | | 0.365 | |
| Model 6 | 0.009 | -51.9 | -50.5 | 0.0013 | | 0.399 | 0.330 |

**Table S6** Phylogenetic analysis results for ETR/J for C3 and C4 species.

| Model | AICw | AIC | AICc | Variance | | Mean | |
|---|---|---|---|---|---|---|---|
| | | | | $C_3$ | $C_4$ | $C_3$ | $C_4$ |
| Model 1 | 0.000 | -16.61 | -16.23 | 0.001 | | 0.825 | |
| Model 2 | 0.000 | -43.71 | -42.94 | 0.005 | 8e-05 | 0.717 | |
| Model 3 | 0.000 | -14.94 | -14.17 | 0.001 | | 0.850 | 0.815 |
| Model 4 | 0.000 | -41.72 | -40.38 | 0.005 | 8e-05 | 0.716 | 0.717 |
| Model 5 | 0.000 | -12.60 | -11.82 | 0.003 | | 0.817 | |
| **Model 6** | **1.000** | **-84.01** | **-82.68** | **0.004** | | **1.082** | **0.643** |

**Table S7** Phylogenetic analysis results for chlorophyll a/chlorophyll b for C3 and C4 species.

| Model | AICw | AIC | AICc | Variance | | Mean | |
|---|---|---|---|---|---|---|---|
| | | | | $C_3$ | $C_4$ | $C_3$ | $C_4$ |
| Model 1 | 0.000 | 44.3 | 44.7 | 0.008 | | 4.004 | |
| Model 2 | 0.002 | 40.8 | 41.6 | 0.018 | 0.003 | 4.178 | |
| Model 3 | 0.000 | 43.6 | 44.4 | 0.008 | | 3.842 | 4.066 |
| Model 4 | 0.003 | 40.1 | 41.5 | 0.018 | 0.003 | 4.054 | 4.224 |
| Model 5 | 0.000 | 49.5 | 50.3 | 0.012 | | 3.972 | |
| **Model 6** | **0.995** | **28.3** | **29.7** | **0.008** | | **3.401** | **4.611** |

**Table S8** Phylogenetic analysis results for ETR for C3 and C4 species.

| Model | AICw | AIC | AICc | Variance | | Mean | |
|---|---|---|---|---|---|---|---|
| | | | | $C_3$ | $C_4$ | $C_3$ | $C_4$ |
| **Model 1** | **0.397** | **333.6** | **334.0** | **30.844** | | **88.941** | |
| Model 2 | 0.165 | 335.4 | 336.1 | 37.380 | 27.334 | 89.019 | |
| Model 3 | 0.148 | 335.6 | 336.4 | 30.798 | | 87.900 | 89.327 |
| Model 4 | 0.063 | 337.3 | 338.6 | 38.118 | 26.881 | 87.284 | 89.643 |
| Model 5 | 0.137 | 335.7 | 336.5 | 85.615 | | 87.068 | |
| Model 6 | 0.091 | 336.5 | 337.9 | 75.383 | | 95.639 | 78.986 |

**Table S9** Phylogenetic analysis results of evolutionary model fitting for Vpmax for C4 species.

| Model | AICw | AIC | AICc | Variance | | Mean | |
|---|---|---|---|---|---|---|---|
| | | | | $C_3$ | $C_4$ | $C_3$ | $C_4$ |
| Model 1 | 0.146 | 213.05 | 213.57 | 7.585 | | 41.583 | |
| **Model 4** | 0.854 | 209.53 | 210.62 | 43.126 | | 41.520 | |

**Table S10** Phylogenetic analysis results of evolutionary model fitting for Vpmax/Vcmax for C4, C3.

| Model | AICw | AIC | AICc | Variance | | Mean | |
|---|---|---|---|---|---|---|---|
| | | | | $C_3$ | $C_4$ | $C_3$ | $C_4$ |
| Model 1 | 0.125 | 6.813 | 7.333 | 0.003 | | 1.908 | |
| **Model 4** | 0.875 | 2.913 | 4.004 | 0.015 | | 1.966 | |

**Table S11** Phylogenetic analysis results for Nitrogen for C3 and C4 species.

| Model | AICw | AIC | AICc | Variance | | Mean | |
|---|---|---|---|---|---|---|---|
| | | | | $C_3$ | $C_4$ | $C_3$ | $C_4$ |
| **Model 1** | 0.184 | 74.82 | 75.20 | 0.013 | | 3.029 | |
| Model 2 | 0.075 | 76.61 | 77.41 | 0.016 | 0.016 | 2.981 | |
| Model 3 | 0.088 | 76.29 | 77.09 | 0.013 | | 2.907 | 3.082 |
| Model 4 | 0.053 | 77.33 | 78.71 | 0.021 | 0.008 | 2.712 | 3.011 |
| Model 5 | 0.037 | 78.01 | 78.81 | 0.029 | | 3.120 | |
| Model 6 | 0.562 | 72.59 | 73.97 | 0.021 | | 3.624 | 2.530 |
